# Female mice exposed to low-doses of dioxin during pregnancy and lactation have increased susceptibility to diet-induced obesity and diabetes

**DOI:** 10.1101/2020.07.09.195552

**Authors:** Myriam P Hoyeck, Rayanna C Merhi, Hannah Blair, C Duncan Spencer, Mikayla A Payant, Diana I Martin Alfonso, Melody Zhang, Geronimo Matteo, Melissa J Chee, Jennifer E Bruin

## Abstract

**Objective:** Exposure to persistent organic pollutants is consistently associated with increased diabetes risk in humans. We investigated the short- and long-term impact of chronic low-dose dioxin (aka 2,3,7,8-tetrachlorodibenzo-*p*-dioxin, TCDD) exposure during pregnancy and lactation on glucose homeostasis and beta cell function in female mice, including their response to a metabolic stressor later in life.

**Methods:** Female mice were injected with either corn oil (CO; vehicle control) or 20 ng/kg/d TCDD 2x/week throughout mating, pregnancy, and lactation, and then tracked for 6-10 weeks after chemical exposure stopped. A subset of CO- and TCDD-exposed dams were then transferred to a 45% high fat diet (HFD) or remained on standard chow diet for an additional 11 weeks to assess long-term effects of TCDD on adaptability to a metabolic stressor.

**Results:** Dioxin-exposed dams were hypoglycemic at birth but otherwise had normal glucose homeostasis during and post-dioxin exposure. However, dioxin-exposed dams on chow diet were modestly heavier than controls starting 5 weeks after the last dioxin injection, and their weight gain accelerated after transitioning to a HFD. Dioxin-exposed dams also had accelerated onset of hyperglycemia, dysregulated insulin secretion, reduced islet size, increased MAFA^-^ beta cells, and increased proinsulin accumulation following HFD feeding compared to control dams.

**Conclusions:** Our mouse model suggests that chronic low-dose dioxin exposure may be a contributing factor to obesity and diabetes pathogenesis in females.

## 1. Introduction

Diabetes is a global health concern, with incidence rates tripling over the past 20 years [1]. This disease is characterized by chronic hyperglycemia, defects in insulin secretion from pancreatic beta cells, and impaired peripheral insulin action [2]. Type 2 diabetes accounts for 90-95% of all diabetes cases and is strongly associated with genetic and lifestyle determinants such as physical inactivity, poor diet, and obesity. Epidemiological studies consistently report significant associations between exposure to persistent organic pollutant (POPs) and increased diabetes incidence in humans [3,4], yet a causal association remains uncertain.

POPs are typically lipophilic chemicals that resist degradation, leading to widespread environmental dispersion and biomagnification [5]. Dioxins and dioxin-like compounds are a broad class of POPs that induce cellular toxicity through persistent activation of the aryl hydrocarbon receptor (AhR), and upregulation of AhR-target genes including cytochrome P450 *(Cyp)1a1* and *Cyp1a2* [6]. CYP enzymes are phase I xenobiotic metabolism enzymes that play an essential role in the oxidation of xenobiotic compounds for subsequent excretion from the body. However, this process generates highly reactive intermediate metabolites which could potentially cause oxidative stress and DNA/protein damage [7]. We previously showed that CYP1A1 is induced and functional in human and mouse pancreatic islets following exposure to the highly persistent dioxin, 2,3,7,8-tetrachlorodibenzo-*p*-dioxin (TCDD, also referred to as “dioxin”) *in vitro* [8]. CYP1A1 was also upregulated in mouse islets after systemic TCDD administration *in vivo* [8,9], indicating that dioxins reach the endocrine pancreas, which may impact beta cell function and survival. Indeed, TCDD-exposed mouse and human islets had significantly reduced glucose-induced insulin secretion *in vitro* [8]. In addition, a single high-dose injection of TCDD (20 µg/kg) *in vivo* reduced plasma insulin levels for up to 6 weeks in male and female mice [9]. TCDD-exposed males also displayed modest fasting hypoglycemia for ∼4 weeks post-injection, increased insulin sensitivity, and decreased beta cell area. In contrast, TCDD-exposed females became transiently glucose intolerant 4 weeks after the single high-dose TCDD injection [9]. These results suggest that pollutants may increase diabetes risk by altering beta cell function and/or islet composition in a sex-dependent manner.

Pregnancy is a unique period of metabolic plasticity in females, during which beta cells proliferate and increase insulin secretion to accommodate for changes in nutritional needs [10–13]. Failure to compensate for these changes can lead to hyperglycemia, gestational diabetes mellitus (GDM), and long-term metabolic complications in mothers and their offspring [14]. Most epidemiological studies investigating the association between POPs and diabetes have focused on adult populations subjected to acute dioxin exposure (e.g. war veterans, occupational workers, and victims of chemical disasters) [15–18], or populations with chronic high background exposure (e.g. populations with high fish consumption) [19–21]. The few studies that assessed the link between chronic low-dose POP exposure during pregnancy and gestational diabetes mellitus (GDM) report inconsistent findings of either a positive association between serum POP levels and GDM [22–25] or no association [25–27]. In addition, these are all cross-sectional studies that do not provide information about the timing of events and cannot rule out the possibility of reverse causality (i.e. whether POPs increased GDM risk, or whether GDM increased serum POP accumulation). Lastly, long-term tracking is not possible with cross-sectional epidemiological studies.

The purpose of our study was first to determine whether chronic low-dose TCDD exposure during pregnancy and lactation impacts pancreatic beta cell plasticity, glucose homeostasis, or body weight regulation during this critical period of metabolic stress in female mice (“TCDD injection” window). We then stopped TCDD administration when offspring were weaned and performed metabolic assessments on the dams for an additional 10 weeks to determine whether the transient TCDD exposure during pregnancy/lactation had lasting metabolic consequences (“Post-TCDD injection” window). Finally, at the end of the “Post-TCDD” window, a subset of control and TCDD-exposed dams were transitioned to high fat diet (HFD) feeding for another 11 weeks to assess their ability to adapt to a metabolic stressor (“Metabolic Challenge” windo,j4w).

## 2. Materials and Methods

### 2.1 Animals

C57BL/6 mice, 5-8 week old (Charles River; Raleigh, NC, USA), were maintained on a 12-hour light/dark cycle throughout the study and received *ad libitum* access to standard rodent chow (Harlan Laboratories, Teklad Diet #2018, Madison, WI). All experiments were approved by the Carleton University Animal Care Committee and carried out in accordance with the Canadian Council on Animal Care guidelines. Prior to beginning experimental protocols, animals were randomly assigned to treatment groups and matched for body weight and blood glucose to ensure that these variables were consistent between groups.

#### 2.1.1. Cohort 1 (Supplemental Fig. S1-S2)

As outlined in **Figure 1A**, female mice received subcutaneous injections (s.c.) of corn oil (CO) (25 ml/kg, vehicle control; n = 6) (#C8267-2.5L, Sigma-Aldrich, St Louise, MO, USA) or a low-dose of TCDD (20 ng/kg/d; n = 6) (Sigma Aldrich, # 48599) 2x/week starting one week prior to pairing with male mice and lasting throughout mating and pregnancy. All mice were euthanized at postnatal day 1 (P1). Whole pancreas was harvested and stored in RNAlater (#76106, Qiagen, Hilden, Germany) for quantitative real-time PCR (qPCR) analysis or 4% paraformaldehyde (PFA; #AAJ19943K2, Thermo Fisher Scientific, Waltham, MA, USA) for 24 hrs, followed by long-term storage in 70% EtOH for histological analysis.

**Fig. 1:**
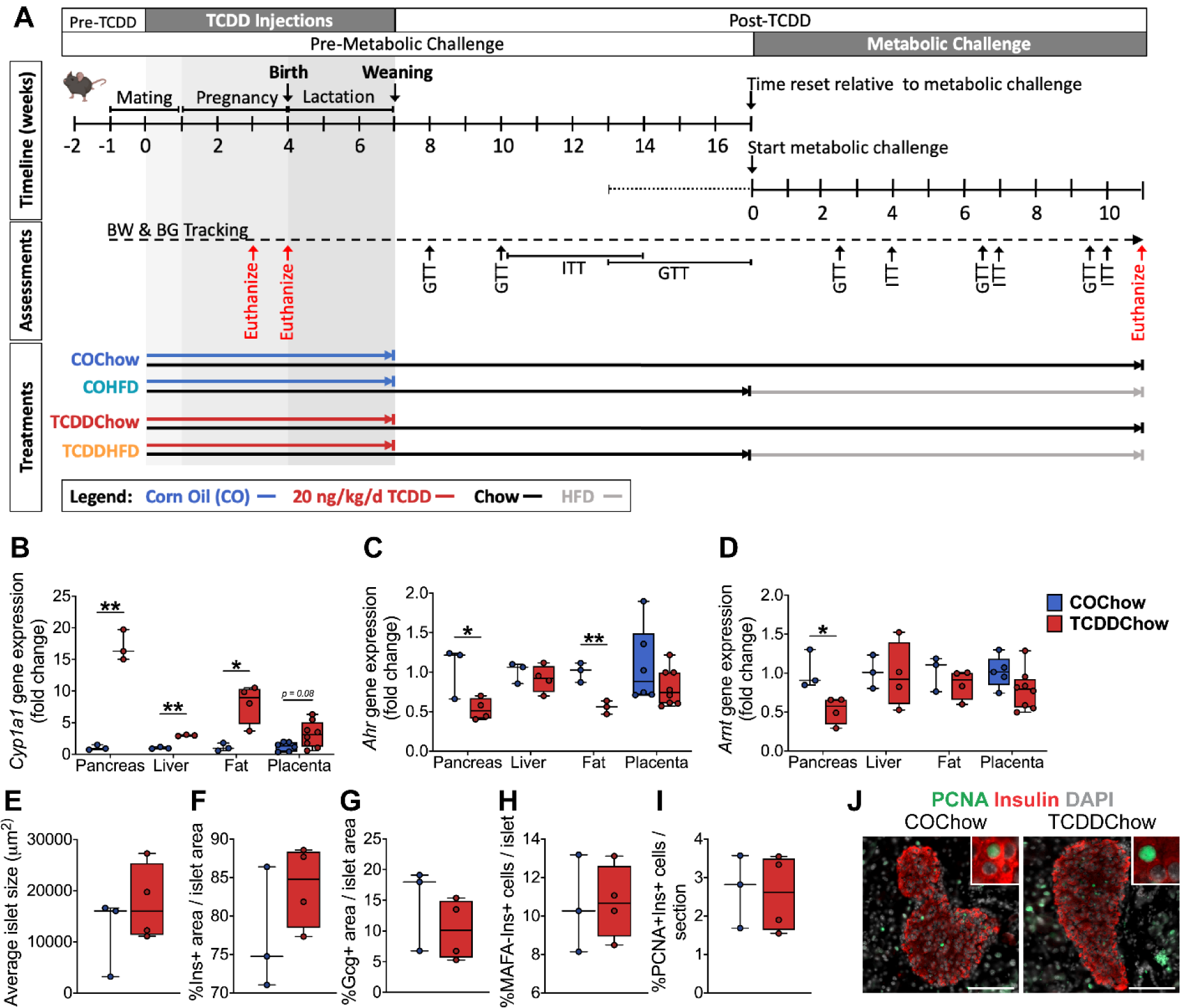
Chronic low-dose TCDD exposure induces *Cyp1a1* expression more strongly in the pancreas than other tissues at mid-gestation. (**A**) Schematic summary timeline of the study. Female mice were injected with either corn oil (CO) or 20 ng/kg/d TCDD 2x/week during mating, pregnancy, and lactation (“TCDD Injections” window), and then tracked for 6-10 weeks after chemical exposure stopped (“Post-TCDD” window) (i.e. week 13-17 of the study). This range reflects differences in the duration of mating, with a 4 week difference between the first and last pregnancies. At weeks 13-17 of the study, a subset of CO- and TCDD-exposed dams were transitioned to a 45% HFD or remained on standard chow. All mice were then assessed for another 11 weeks. BW = body weight; BG = blood glucose; GTT = glucose tolerance test; ITT = insulin tolerance test. (**B-D**) Gene expression was measured in whole pancreas, liver, perirenal fat, and placenta on day 15.5 of pregnancy, including (**B**) *Cyp1a1*, (**C**) *Ahr*, and (**D**) *Arnt* gene expression. Gene levels are expressed as fold change relative to control. (**E-J**) Histological analysis of pancreas collected from dams mid-gestation, including quantification of (**E**) average islet area, (**F**) % insulin^+^ area / islet area, (**G**) % glucagon^+^ area / islet area, (**H**) % MAFA^-^INS^+^ cells per islet, and (**I**) % PCNA^+^INS^+^ cells per pancreas section. (**J**) Representative images of pancreas sections showing immunofluorescent staining for insulin and PCNA. Inset regions show PCNA^+^INS^+^ cells. Scale bar = 100 μm. All data are presented as median with min/max values. Individual data points on box and whisker plots represent biological replicates (different mice). *p<0.05, **p<0.01 versus control. The following statistical tests were used: (**B**) two-tailed unpaired t-test for pancreas, liver, and fat; Mann Whitney Test for placenta; (**C-D, E-I**) two-tailed unpaired t-test.

#### 2.1.2. Cohort 2 (Figure 1-6; Supplemental Fig. S3-S4)

As outlined in **Figure 1A**, female mice received s.c. injections of CO (25 ml/kg, vehicle control; n = 12) or a low-dose of TCDD (20 ng/kg/d; n = 14) 2x/week starting one week prior to pairing with male mice and lasting throughout mating, pregnancy, and lactation (“TCDD Injections” window: weeks 0-7 of the study). Metabolic assessments were performed on chow-fed dams for 6 to 10 weeks following the last TCDD exposure at weaning (“Post-TCDD” window: weeks 7-17 of the study). This range reflects differences in the duration of mating, with a 4-week difference between the first and last pregnancies. As outlined in **Figure 1A**, a subset of CO- and TCDD-exposed dams were then transferred to a 45% HFD (Research Diets D12451, New Brunswick, NJ) or remained on a standard chow diet for an additional 11 weeks (“Metabolic Challenge” window), generating the following experimental groups (n=4-5 per group): COChow, COHFD, TCDDChow, and TCDDHFD. Whole pancreas, liver, perirenal fat, and placenta were harvested from a subset of mice at approximately day 15.5 of pregnancy (GD15.5) (n=3-4/group). Hypothalamic brain tissue was collected at week 11 of the metabolic challenge (n=4-5/group). Tissues were stored in RNAlater or 4% PFA.

#### 2.1.3. Cohort 3 (Figure 7)

As outlined in **Figure 7A**, non-pregnant female mice received intraperitoneal (i.p.) injections of CO (25 ml/kg, vehicle control) or a low-dose of TCDD (20 ng/kg/d) 2x/week for 12 weeks. During this period of active chemical exposure, a subset of CO- and TCDD-exposed females were fed either standard chow diet or 45% HFD, generating the following experimental groups (n=3-4/group): COChow, COHFD, TCDDChow, and TCDDHFD. After 12 weeks of exposure, pancreatic islets were isolated from all mice and stored in RNAlater for subsequent analysis by RNAseq and qPCR.

**Fig. 2:**
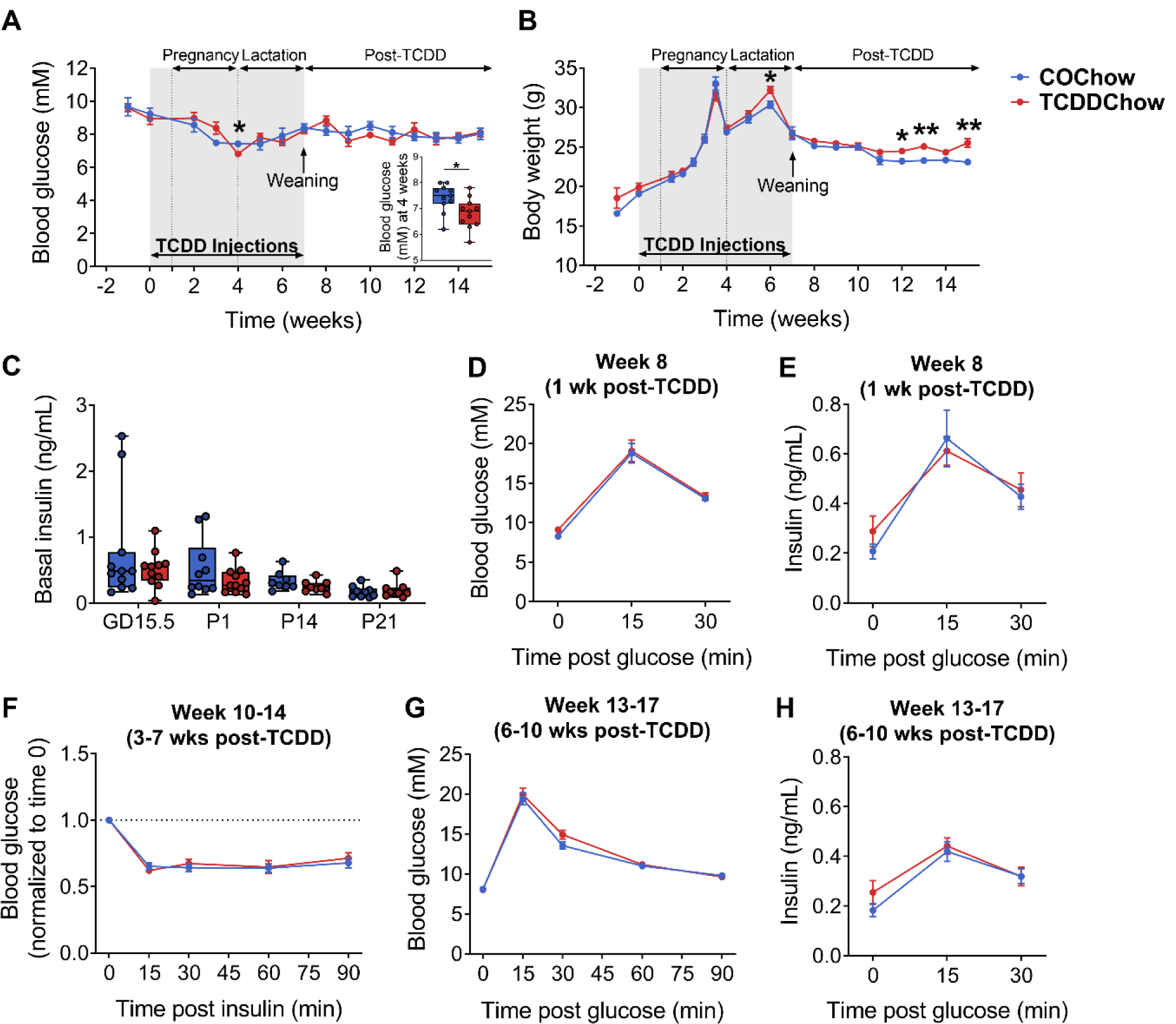
Transient TCDD exposure during pregnancy and lactation causes hypoglycemia at birth and promotes weight gain post-exposure. Female mice were injected with either corn oil (CO) or 20 ng/kg/d TCDD 2x/week during mating, pregnancy, and lactation, and then tracked for 6-10 weeks after exposure (see **Figure 1A** for study timeline). GTT = glucose tolerance test; ITT = insulin tolerance test. (**A**) Blood glucose and (**B**) body weight were measured weekly following a 4-hour morning fast. Blood glucose at postnatal day 1 (P1) is shown in the inset graph in (**A**). (**C**) Plasma insulin levels were measured at day 15.5 of pregnancy, P1, mid-lactation (P14), and weaning (P21) following a 4-hour morning fast. (**D, G**) Blood glucose and (**E, H**) plasma insulin levels during a GTT at (**D, E**) 1 week post-TCDD (i.e. week 8 of the study) and (**G, H**) 6-10 weeks post-TCDD (week 13-17 of the study). (**F**) Blood glucose levels during an ITT at 3-7 weeks post-TCDD (week 10-14 of the study) (values are normalized relative to time zero for each mouse). All data are presented as mean ± SEM in line graphs or median with min/max values in box and whisker plots. Individual data points on box and whisker plots represent biological replicates (different mice). *p <0.05, **p <0.01 versus control. The following statistical tests were used: (**A, B**) line graphs, two-way REML-ANOVA with uncorrected Fisher’s LSD test; box and whisker plots, two-tailed unpaired t-test; (**C**) two-tailed unpaired t-test at P14; two-tailed Mann-Whitney test at GD15, P0, and P12; (**D-H**) two-way RM ANOVA with Sidak test.

**Fig. 3:**
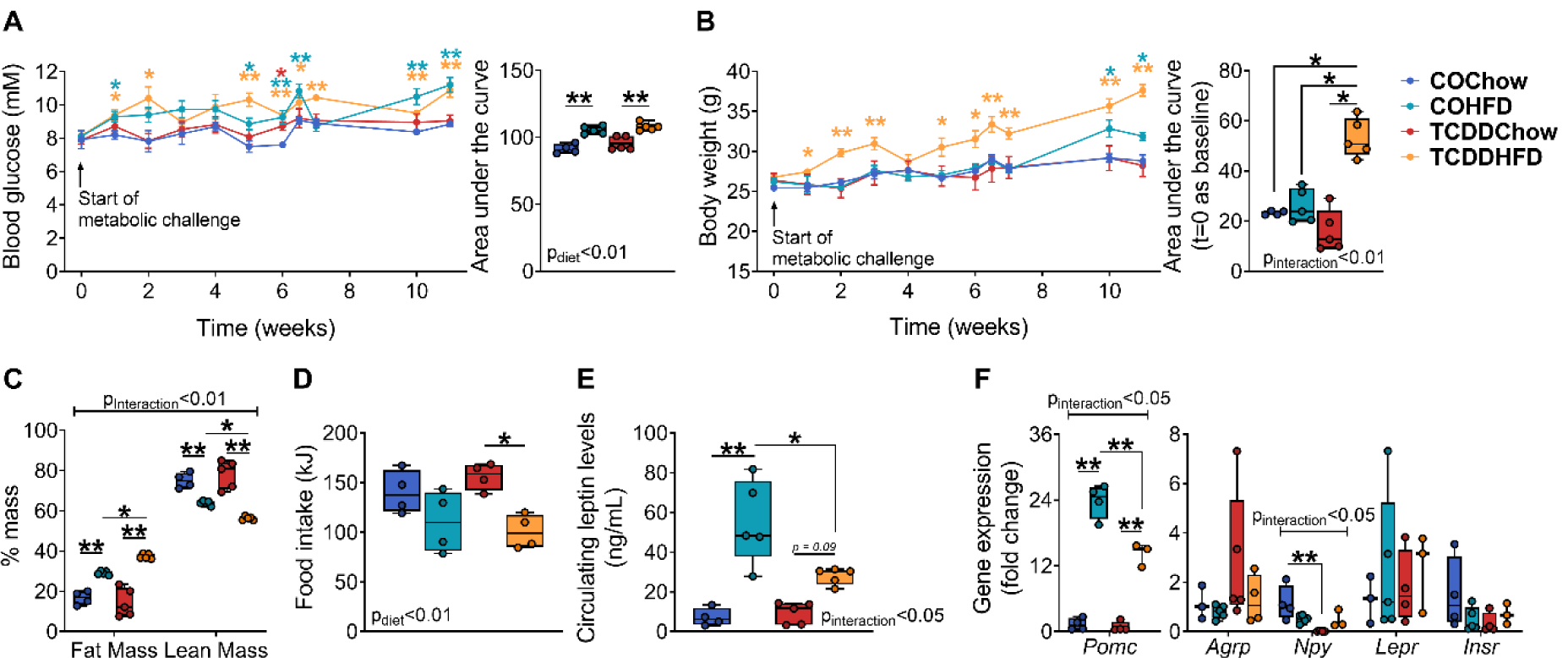
TCDD exposure promotes diet-induced obesity in dams and alters energy homeostasis. Following 6-10 weeks of post-TCDD tracking in chow-fed mice (i.e. weeks 13-17 of the study), a subset of CO- and TCDD-exposed dams were transitioned to a 45% HFD or remained on standard chow and assessed for another 11 weeks (“Metabolic Challenge” window) (see **Figure 1A** for study timeline). (**A**) Blood glucose and (**B**) body weight were measured weekly following a 4-hour morning fast. (**C**) % fat mass and lean mass were measured by EchoMRI 10 weeks into the metabolic challenge. (**D**) Caloric intake was measured at week 7 of the challenge. (**E**) Circulating leptin levels were measured in cardiac blood of random-fed dams at week 11 of the challenge. (**F**) Gene expression for markers of energy homeostasis were measured in the hypothalamus at week 11 of the challenge. All data are presented as mean ± SEM in line graphs or median with min/max values in box and whisker plots. Individual data points on box and whisker plots represent biological replicates (different mice). *p <0.05, **p<0.01, coloured stars are versus COChow. The following statistical tests were used: (**A, B**) line graphs, two-way RM ANOVA with uncorrected Fisher’s LSD test; box and whisker plots, two-way ANOVA with Tukey’s multiple comparison test; (**C-F**) two-way ANOVA with Tukey’s multiple comparison test.

**Fig. 4:**
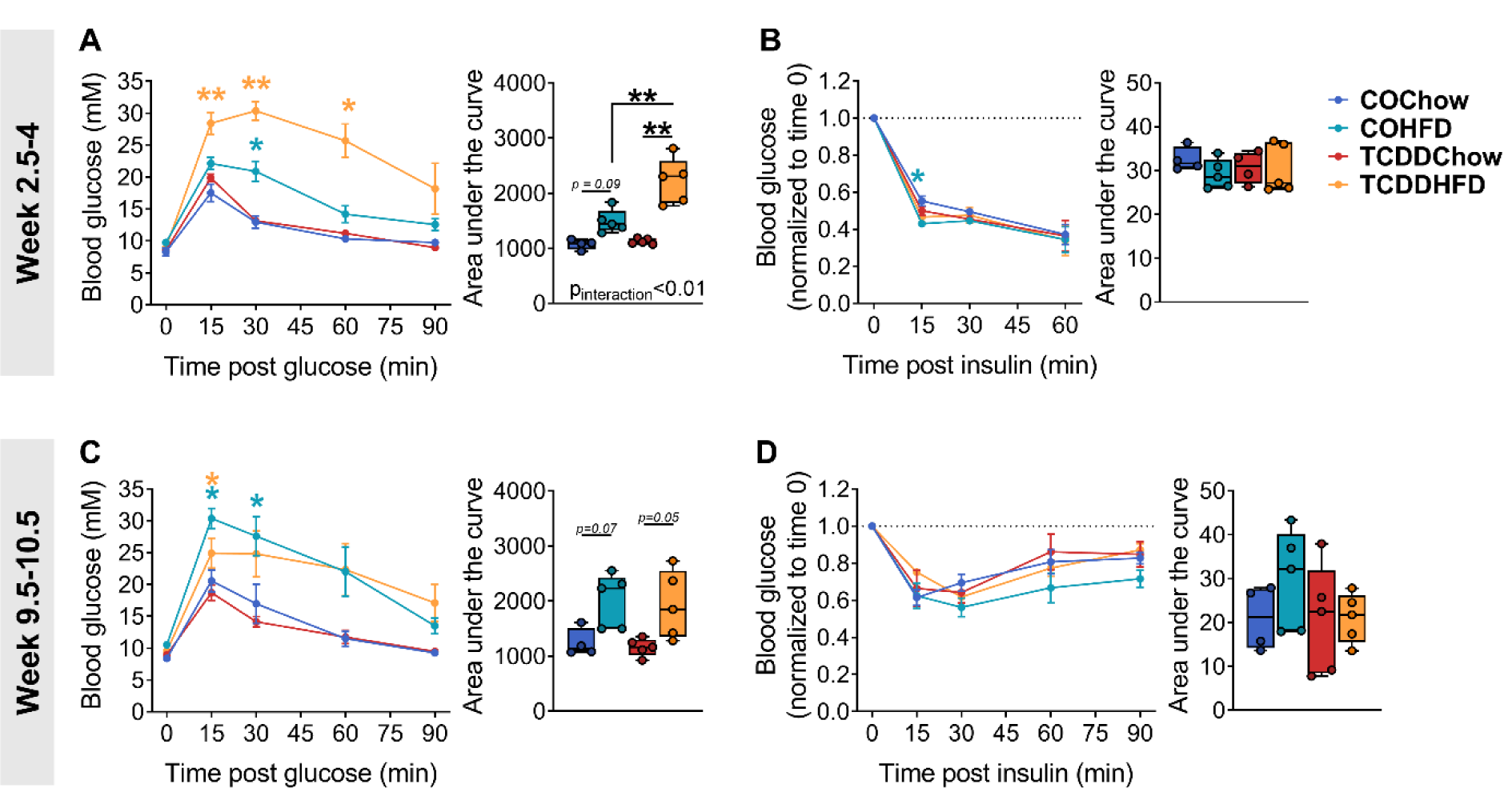
HFD feeding causes severe hyperglycemia in TCDD-exposed dams without altering insulin tolerance. Glucose and insulin tolerance were assessed *in vivo* at weeks 2.5-4 and 9.5-10.5 of the metabolic challenge (see **Figure 1A** for study timeline). (**A, C**) Blood glucose levels during a glucose-tolerance test at (**A**) week 2.5 and (**C**) week 9.5 of the metabolic challenge. (**B, D**) Blood glucose levels during an insulin tolerance test at (**B**) week 4 and (**D**) week 10.5 of the metabolic challenge (values are normalized relative to time zero for each mouse). All data are presented as mean ± SEM in line graphs or median with min/max values in box and whisker plots. Individual data points on box and whisker plots represent biological replicates (different mice). *p <0.05, **p <0.01, coloured stars are versus COChow. The following statistical tests were used: (**A-D**) line graphs, two-way RM ANOVA with Tukey’s multiple comparison test; box and whisker plots, two-way ANOVA with Tukey’s multiple comparison test.

**Fig. 5:**
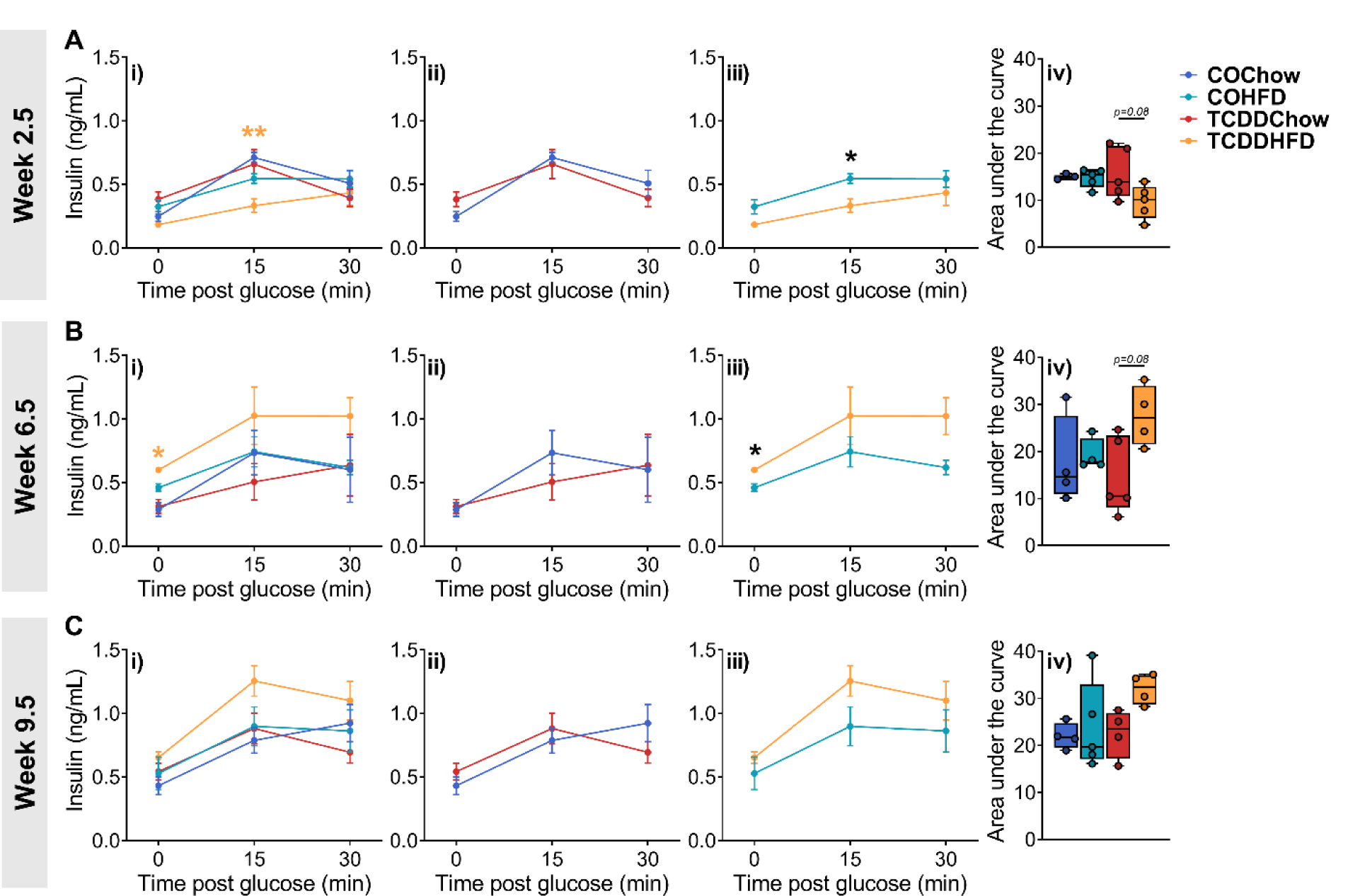
HFD feeding impairs insulin secretion in TCDD-exposed dams. Glucose-stimulated insulin secretion was assessed *in vivo* at (**A**) week 2.5, (**B**) week 6.5, and (**C**) week 9.5 of the metabolic challenge (see **Figure 1A** for study timeline). Insulin data are presented as (**i**) all groups compared to COChow, (**ii**) TCDDChow versus COChow, (**iii**) TCDDHFD compared to COHFD, and (**iv**) area under the curve for all groups. All data are presented as mean ± SEM in line graphs or median with min/max values in box and whisker plots. Individual data points on box and whisker plots represent biological replicates (different mice). *p <0.05, coloured stars are versus COChow. The following statistical tests were used: (**A-C i**) two-way RM ANOVA with Tukey’s multiple comparison test; (**A-C ii, iii**) two-way RM ANOVA with Sidak test; (**A-C iv**) two-way ANOVA with Tukey’s multiple comparison test.

**Fig. 6:**
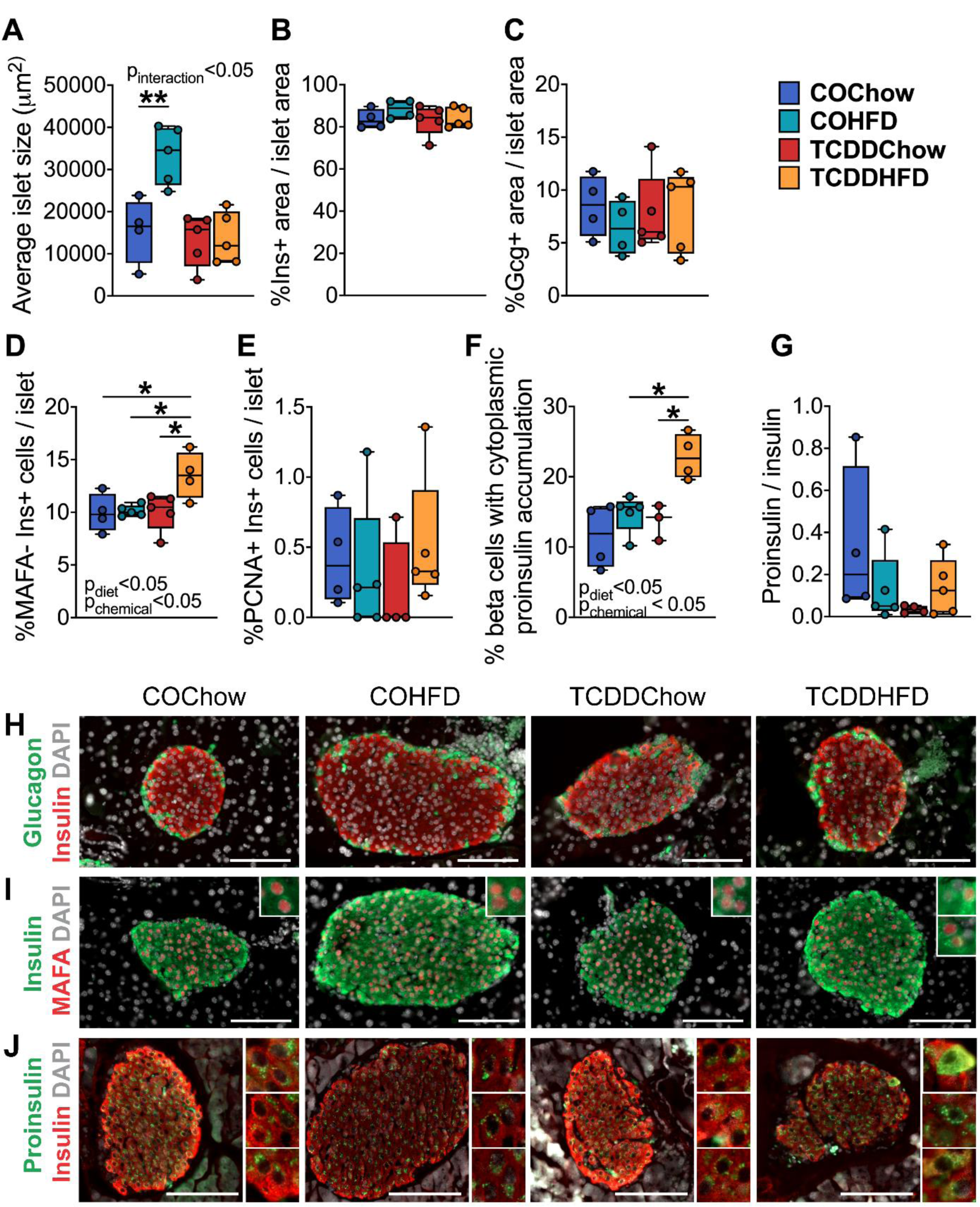
TCDDHFD dams have decreased islet size, a higher proportion of MAFA^-^ beta cells, and increased proinsulin accumulation compared to COHFD dams. Whole pancreas and cardiac blood were harvested at week 11 of the metabolic challenge for analysis (see **Figure 1A** for study timeline). (**A**) Average islet area, (**B**) % islet^+^ area / islet area, (**C**) % glucagon^+^ area / islet area, (**D**) MAFA^-^INS^+^ cells per islet, (**E**) % PCNA^+^INS^+^ cells per islet, and (**F**) % beta cells with cytoplasmic proinsulin accumulation, as determined by immunofluorescent staining. (**G**) Circulating proinsulin/insulin levels in cardiac blood of random-fed dams. (**H-J**) Representative images of pancreas sections showing immunofluorescence staining for (**H**) insulin/glucagon, (**I**) insulin/MAFA, or (**J**) insulin and proinsulin. Inset regions in **(I)** show MAFA^+^INS^+^ cells. Inset regions in (**J**) show perinuclear proinsulin immunoreactivity versus proinsulin accumulation in the cytoplasm. Scale bar = 100 µm. All data are presented as median with min/max values. Individual data points on box and whisker plots represent biological replicates (different mice). *p <0.05, **p <0.01. The following statistical tests were used: (**A-G**) two-way ANOVA with Tukey’s multiple comparison test.

**Fig. 7:**
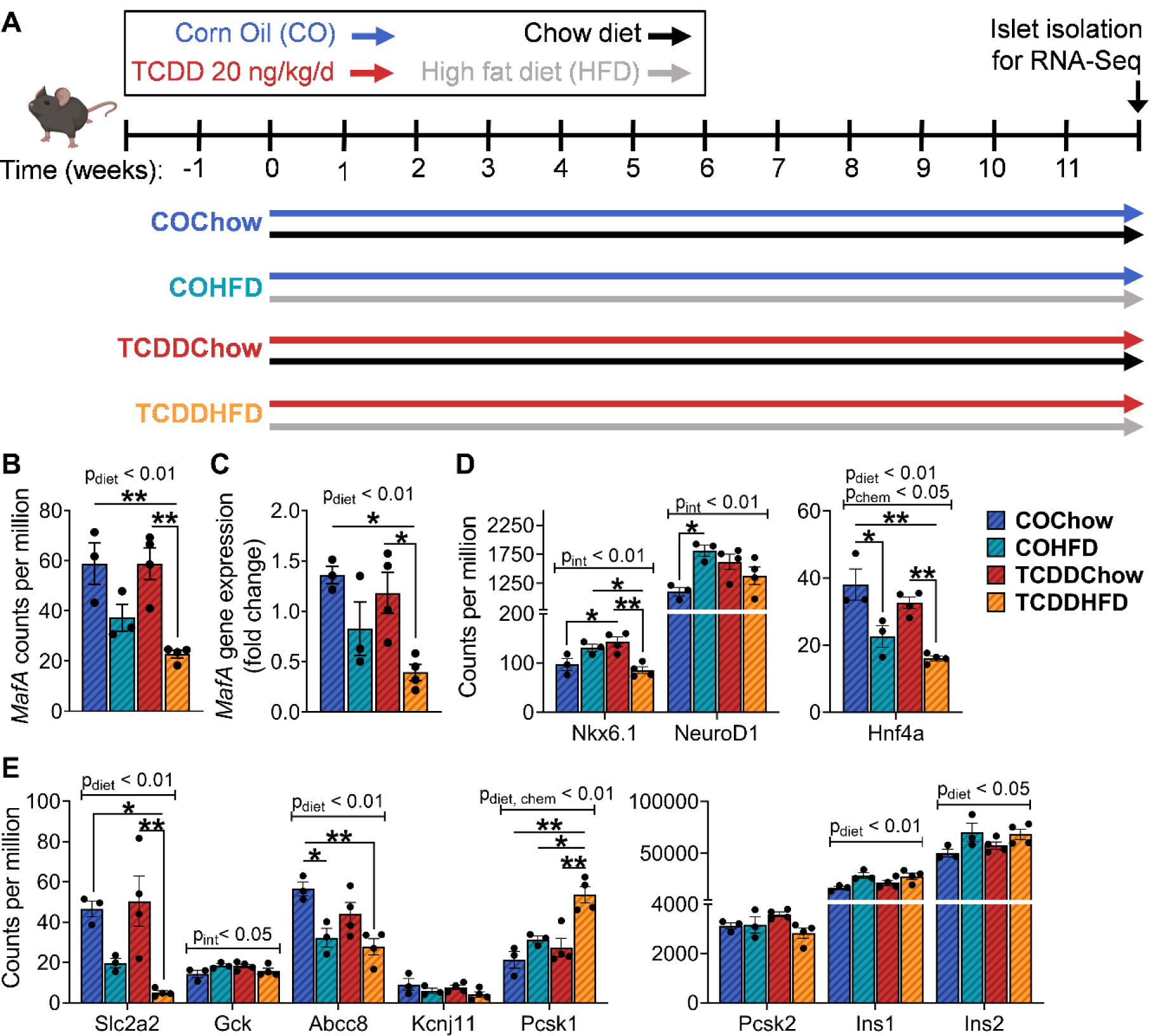
TCDDHFD females have reduced *MafA, Nkx6*.*1* and *Slc2a2* gene expression, and increased *Pcsk1* expression compared to TCDDChow females. (**A**) Schematic summary timeline of the study. Female mice were injected with corn oil (CO) or 20 ng/kg/d TCDD 2x/week for 12 weeks, and either fed standard chow or 45% HFD. Islets were isolated at 12 weeks for subsequent gene expression analysis by RNAseq and qPCR. (**B, D-E**) RNAseq data showing gene counts per million for (**B**) MafA, (**D**) beta cell transcription factors, and (**E**) genes associated with insulin processing and secretion. (**C**) qPCR analysis of MafA expression. All data are presented as mean ± SEM. Individual data points on bar graphs represent biological replicates (different mice). *p <0.05, **p <0.01. The following statistical tests were used: (**B-E**) two-way ANOVA with Tukey’s multiple comparison test.

### 2.2. Metabolic Assessment

All metabolic analyses were performed in conscious, restrained mice, and blood samples were collected via the saphenous vein using heparinized microhematocrit tubes. Blood glucose levels were measured using a handheld glucometer (Lifescan; Burnaby; Canada)

Body weight and blood glucose were measured weekly or 2x/week following a 4-hour morning fast. Saphenous blood was collected at GD15.5, P1, lactation (P14), and weaning (P21) following a 4-hour morning fast to measure plasma insulin levels by ELISA (#80-INSMSU-E01, ALPCO, Salem, NH, USA). For all metabolic tests, time 0 indicates the blood sample collected prior to administration of glucose or insulin. For glucose tolerance tests (GTTs), mice received an i.p. bolus of glucose (2 g/kg) following a 4-hour morning fast. Blood samples were collected at 0, 15, and 30 minutes for measuring plasma insulin levels by ELISA. For insulin tolerance tests (ITTs), mice received an i.p. bolus of insulin (0.7 IU/kg: **Figure 2F, 4D** and **Supplemental Fig. S4B**; 1.0 IU/kg: **Figure 4B**) (Novolin ge Toronto; Novo Nordisk Canada, #02024233) following a 4-hour morning fast. Cardiac blood was collected at week 11 of the metabolic challenge for measuring random-fed plasma leptin levels (#90030, Crystal Chem, Elk Grove Village, IL, USA), proinsulin levels (ALPCO, #80-PINMS-E01), and insulin levels (ALPCO, #80-INSMSU-E01) by ELISA.

### 2.3. Food Intake Analysis

Food intake was assessed mid-pregnancy, lactation (P14), and at week 7 of the metabolic challenge. All dams were singly housed for this experiment. Dams were transferred to a clean cage and the amount of food in their feeding trays was weighed (Day 0). Food weight was measured again 2 days later (Day 2), ensuring that any large pieces of food that had fallen into the cage were included in the measurement. Food intake in kcal was determined as follows: [(Weight_Day 0_ – Weight_Day 2_) x Food Energy Density], where standard chow has an energy density of 13.0 kJ/g and 45% HFD a density of 19.7 kJ/g.

### 2.4. EchoMRI Analysis

Fat mass and lean mass were measured by the University of Ottawa Animal Behaviour Core (Ottawa, ON) at week 10 of the metabolic challenge using an EchoMRI-700 (EchoMRI LLC, Houston, TX, USA). Percent fat and lean mass were determined relative to total body weight measured immediately prior to doing the MRI. The instrument was calibrated as per the manufacturer’s instructions prior to measurements.

### 2.5. Islet isolation and culture

Islets were isolated by pancreatic duct injection with collagenase (1000 U/ml; Sigma-Aldrich, #C7657) as previously described [8]. In brief, inflated pancreas tissues were excised and incubated at 37°C for 12 min, vigorously agitated, and the collagenase reaction quenched. The pancreas tissue was washed three times in HBSS+CaCl_2_ and resuspended in Ham’s F-10 (#SH30025.01, HyClone, GE Healthcare Bio-sciences, Pittsburgh, PA, USA). Pancreas tissue was filtered through a 70 μm cell strainer and islets were handpicked under a dissecting scope to >95% purity.

### 2.6. Quantitative Real Time PCR

RNA was isolated from whole pancreas, liver, perirenal fat, placenta, and hypothalamic brain using TRIzol™ (#15596018, Invitrogen, Carlsbad, CA, USA) as per the manufacturer’s instructions. RNA was isolated from islets using the RNeasy Micro Kit (Qiagen, #74004) as per the manufacturer’s instructions, with the following amendment: 7 volumes of buffer RLT + DTT were added to the samples prior to lysing with 70% EtOH. DNase treatment was performed prior to cDNA synthesis with the iScript™ gDNA Clear cDNA Synthesis Kit (#1725035, Biorad, Mississauga, ON, Canada). qPCR was performed using SsoAdvanced Universal SYBR Green Supermix (Biorad, #1725271) or SensiFAST™ SYBR^®^ No-ROX Supermix (# BIO-980050, Bioline, London, UK) and run on a CFX96 or CFX394 (Biorad). *Hprt, Ppia*, or *CypA* was used as the reference gene since these genes displayed stable expression under control and treatment conditions. Data were analyzed using the 2^ΔΔCT^ relative quantitation method. Primer sequences are listed in **Table S1**.

### 2.7. RNA Sequencing

Gene expression in whole islets was measured using the TempO-Seq Mouse Whole Transcriptome panel (BioSpyder Technologies Inc, Carlsbad, CA). 2 µl of RNA was diluted in 2x TempO-Seq lysis buffer diluted with an equal amount of PBS. Lysates, positive controls (100 ng/µl of Universal Mouse Reference RNA (UMRR) Quantitative PCR Mouse Reference Total RNA Agilent, cat # 750600), and no-cell negative controls (1X TempO-Seq lysis buffer alone) were hybridized to the detector oligo mix following the manufacturer’s instructions (TEMPO-SEQ Mouse Mouse Whole Transcriptome AssayTranscriptome Kit (96 Samples) BioSpyder Technologies, Inc. Carlsbad, CA, USA). Hybridization was followed by nuclease digestion of excess oligos, detector oligo ligation, and amplification of the product with the tagged primers according to manufacturer’s instructions. Each sample’s primers were also ligated to a sample-specific barcode that allows multiplexing for sequencing purposes. Labelled amplicons were pooled and purified using NucleoSpin Gel and PCR Clean-up kits (Takara Bio USA, Inc, Mountain View, CA USA). Libraries were sequenced in-house using a NextSeq 500 High-Throughput Sequencing System (Illumina, San Diego, CA, USA) using 50 cycles from a 75-cycle high throughput flow cell. A median read depth of 2 million reads/sample was achieved.

Reads were extracted from the bcl files, demultiplexed (i.e., assigned to respective sample files) and were processed into fastq files with bcl2fastq v.2.17.1.14. The fastq files were processed with the “pete. star. script_v3.0” supplied by BioSpyder. Briefly, the script uses star v.2.5 to align the reads and the qCount function from QuasR to extract the feature counts specified in a gtf file from the aligned reads. The data were then passed through internal quality control scripts. Boxplots of the log2 CPM (counts per million) were plotted to ensure a reproducible distribution between replicates within a group. Hierarchical clustering plots were generated (hclust function: default linkage function of hclust function in R; complete-lin kage) using a distance metric defined as 1-Spearman correlation in order to identify potential outliers. Probes with low counts (i.e. less than a median of 5 counts in at least one group) were flagged as absent genes and eliminated from the dataset. DEG analysis was conducted using the R software [28] on the counts using the default parameters of DESeq2 [29] with respective control and exposure groups. A shrinkage estimator was applied to the fold change estimates using the apeglm method [30] using the lfcShrink() function.

### 2.8 Immunofluorescent Staining and Image Quantification

Whole pancreas tissues were processed and paraffin-embedded by the University of Ottawa Heart Institute Histology Core Facility (Ottawa, ON). Tissues were sectioned (5 µm thick) using a Thermo Scientific Microm HM 355S. Immunofluorescent staining was performed as previously described [8]. In brief, slides were deparaffinized with sequential incubations in xylene and ethanol. Heat-induced epitope retrieval was performed in 10 mM citrate buffer or 10 mM Tris base at 95°C for 10-15 minutes, and slides were incubated with Dako Serum Free Protein Block (#X090930-2, Agilent, Santa Clara, CA) for 30 minutes. Slides were incubated overnight at 4°C with primary antibodies, and then incubated with secondary antibodies for 1-hour at room temperature. Coverslips were mounted with Vectashield® hardset mounting medium with DAPI (#H-1500, Vector Laboratories, Burlingame, CA, USA) or ProLong Gold antifade mountant (Thermo Fisher Scientific, #P36935) for counterstaining nuclei.

The following primary antibodies were used: rabbit anti-insulin (1:200, C27C9, Cell Signaling Technology, Danvers, MA, USA; #3014), mouse anti-insulin (1:250, L6B10, Cell Signaling, #8138S), mouse anti-glucagon (1:250, Sigma-Aldrich, #G2654), mouse anti-PCNA (1:100, BD Transduction Laboratories, San Jose, CA, USA; #610665), rabbit anti-MAFA (1:1000, shared by Dr. Timothy Kieffer), and mouse anti-proinsulin (1:50, DSHB, Iowa City, IA, USA; GS-9A8-s). The following secondary antibodies were used: goat anti-rabbit IgG (H+L) secondary antibody, Alexa Fluor 594 (1:1000, Invitrogen, #A11037); and goat anti-mouse IgG (H+L) secondary antibody, Alexa Fluor 488 (1:1000, Invitrogen, #A11029).

For islet morphology quantification, a range of 5-13 islets were imaged per mouse with an Axio Observer 7 microscope and the average of all islet measurements is reported for each biological replicate. Immunofluorescence was manually quantified using Zen Blue 2.6 software (Carl Zeiss, Germany). The % hormone^+^ area per islet was calculated as [(hormone^+^ area / islet area) x 100]. The % PCNA^+^ insulin^+^ cells per islet was calculated as [(# of PCNA^+^ insulin^+^ cells) / (total # of insulin^+^ cells per islet) x 100], with an average of 590 cells counted per mouse. The % MAFA^-^ insulin^+^ cells per islet was calculated as [(# of MAFA^-^ insulin^+^ cells per islet) / (total # of insulin^+^ cells per islet) x 100], with an average of 750 cells counted per mouse. The % beta cells with accumulation proinsulin was calculated as [(# of insulin^+^ cells with accumulated cytoplasmic proinsulin) / (total # of insulin^+^ cells per islet) x 100], with an average of 400 cells counted per mouse.

### 2.9. Quantification and Statistical Analysis

All statistics were performed using GraphPad Prism 8.4.2 (GraphPad Software Inc., La Jolla, CA). Specific statistical tests are indicated in figure legends and summarized in **Table S2**. Verification of parametric assumptions are found in **Table S2**. Sample sizes are described in section **2.1 Animals**. For all analyses, p<0.05 was considered statistically significant. Statistically significant outliers were detected by a Grubbs’ test with α=0.05. All data was tested for normality using a Shapiro-Wilk test and for equal variance using either a Brown-Forsyth test (for one-way ANOVAs) or an F test (for unpaired t tests). Non-parametric statistics were used in cases where the data failed normality or equal variance tests. Parametric tests were used for all two-way ANOVAs, but normality and equal variance were tested on area under the curve values and by one-way ANOVAs (**Table S2**). A two-way mixed-effect model (REML) ANOVA was used in cases where data points were missing due to random reasons. Box and whisker plots display the median, with whiskers representing minimum to maximum values. Bar plots display mean ± SEM. Individual data points on box and whiskers or bar plots are always biological replicates (i.e. different mice).

### 2.10. Data and Resource Availability

The datasets generated and/or analyzed during the current study are available from the corresponding author upon reasonable request.

## 3. Results

### 3.1. Chronic low-dose TCDD exposure induces Cyp1a1 gene expression more strongly in the pancreas than other tissues at mid-gestation

Briefly, female mice were injected with either corn oil (CO; vehicle control) or 20 ng/kg/d TCDD 2x/week during mating, pregnancy, and lactation (“TCDD Injections” window), and then tracked for 6-10 weeks after chemical exposure stopped (“Post-TCDD” window) (**Figure 1A**). A subset of CO- and TCDD-exposed dams were then transferred to a 45% HFD or remained on standard chow diet for an additional 11 weeks to assess long-term effects of TCDD on adaptability to a metabolic stressor (“Metabolic Challenge” window) (**Figure 1A**).

Before examining the effects of chronic low-dose TCDD exposure on glucose homeostasis and beta cell adaptation during pregnancy, we first validated our model by ensuring that TCDD was reaching the dam pancreas mid-gestation using *Cyp1a1* as an indicator of direct pollutant exposure [8]. TCDD induced *Cyp1a1* expression ∼17-fold in the pancreas compared to ∼3-fold, ∼8-fold, and ∼3.5-fold increase in the liver, perirenal fat, and placenta, respectively, at day 15.5 of pregnancy relative to control (**Figure 1B**). *Ahr* expression was reduced ∼2-fold in the pancreas and perirenal fat of TCDD-exposed dams compared to controls, and not changed in liver or placenta (**Figure 1C**). *Arnt* was also reduced ∼2-fold in the pancreas but was not changed in other tissues compared to control (**Figure 1D**). These results indicate that TCDD reaches the pancreas of pregnant dams and strongly induces *Cyp1a1* expression, which leads to negative feedback regulation of the AhR pathway.

We next assessed whether maternal TCDD exposure altered islet morphology or beta cell characteristics mid-pregnancy, a period known to be associated with increased beta cell proliferation [10–13]. TCDD exposure did not alter the average islet area (**Figure 1E, 1J**), % insulin^+^ area per islet (**Figure 1F**), % glucagon^+^ area per islet (**Figure 1G**), % MAFA^-^ beta cells (**Figure 1H**), or % PCNA^+^ beta cells (**Figure 1I-J**). Therefore, chronic low-dose TCDD exposure during pregnancy in mice does not influence beta cell proliferation, beta cell maturity, or islet composition at gestation day 15.5.

### 3.2. TCDD exposure during pregnancy and lactation promotes weight gain post-exposure

We next investigated whether transient TCDD exposure during pregnancy and lactation had short- or long-term effects on dam metabolism. TCDD-exposed dams were transiently hypoglycemic on postnatal day 1 (P1) compared to control dams (**Figure 2A-inset**), but otherwise had normal fasting glycemia during TCDD exposure and for 10 weeks after exposure ended (i.e. Post-TCDD; **Figure 2A**). Importantly, the TCDD-induced hypoglycemia at P1 was replicated in a second independent cohort of mice (**Supplemental Fig. S1A**). There were also no changes in fasting plasma insulin levels during pregnancy or lactation with either cohort (**Figure 2C; Supplemental Fig. S1C**), or in islet morphology of dam pancreas at P1 (**Supplemental Fig. S2A-G)**. Since we did not observe any signs of GDM development in TCDD-exposed dams, we did not perform glucose or insulin tolerance tests during pregnancy to avoid unnecessary stress. However, we did conduct extensive metabolic testing on the dams for 10 weeks after TCDD exposure stopped at the end of lactation. TCDD exposure did not cause any long-term changes to glucose tolerance (**Figure 2D, 2G; Supplemental Fig. S3B**), glucose-stimulated insulin secretion (**Figure 2E, 2H; Supplemental Fig. S3C**), or insulin tolerance (**Figure 2F**) in the 10 weeks post-TCDD (i.e. week 7-17 of the study).

We also assessed the impact of TCDD on body weight during and after exposure. Throughout pregnancy TCDD-exposed dams had normal body weight (**Figure 2B; Supplemental Fig. S1B**) and food intake (**Supplemental Fig. S3A**). However, TCDD-exposed dams were transiently heavier than controls mid-lactation (**Figure 2B**), with no change in food intake (**Supplemental Fig. S3A**). We also observed a modest but statistically significant increase in body weight in TCDD-exposed dams starting 5 weeks after their last TCDD exposure (**Figure 2B**). Based on these results, we speculated that female mice who were exposed to TCDD during pregnancy and lactation might be more susceptible to weight gain when challenged with an obesogenic diet (45% HFD).

### 3.3. Transient TCDD exposure during pregnancy and lactation promotes diet-induced obesity and alters energy homeostasis in dams

Between 6-10 weeks after the last TCDD exposure (week 13-17 of the study), a subset of CO- and TCDD-exposed dams were transitioned to a 45% HFD (COHFD, TCDDHFD) while the other subset remained on standard chow (COChow, TCDDChow) for 11 weeks (see **Figure 1A** for study timeline). Consistent with our findings in **Figure 2A**, TCDDChow dams continued to maintain normal fasting glycemia compared to COChow dams throughout the study (**Figure 3A**). HFD-fed dams developed moderate fasting hyperglycemia, irrespective of chemical exposure (**Figure 3A**). During this “metabolic challenge” window, we were particularly interested in whether TCDD-induced weight gain would persist long-term (i.e. TCDDChow versus COChow), and whether HFD feeding would exacerbate these effects (i.e. TCDDHFD versus TCDDChow). The increased body weight in TCDDChow dams (**Figure 2B**) returned to control levels after longer tracking (**Figure 3B**). Interestingly, TCDDHFD dams showed a rapid and sustained increase in body weight starting 1 week after being transferred to HFD, whereas COHFD dams only had a modest increase in body weight after ∼10 weeks of HFD feeding (**Figure 3B**). Both HFD-fed groups had significantly increased % fat mass and decreased % lean mass compared to chow-fed dams after 10 weeks of HFD feeding, but these changes in body composition were exacerbated in TCDDHFD dams compared to COHFD dams (**Figure 3C**).

To better understand why TCDD-exposed mice gained more fat mass than control mice after HFD feeding, we investigated other aspects of energy homeostasis. We observed a significant decrease in caloric intake in TCDDHFD dams after 7 weeks of HFD feeding compared to TCDDChow dams but this was not different than COHFD dams, so unlikely to explain the dramatic differences in weight gain between HFD groups (**Figure 3D**). As expected, circulating leptin levels were significantly increased ∼7.7-fold in COHFD dams compared to COChow dams after 11 weeks on HFD (**Figure 3E)**. In contrast, TCDDHFD dams only showed a non-significant trend towards having ∼3-fold higher circulating leptin compared to TCDDChow dams, and leptin levels in TCDDHFD dams were significantly lower than COHFD dams (**Figure 3E**). To assess the contribution of central energy balance in TCDD-mediated obesity, we measured gene expression of key regulators within the brain. Proopiomelanocotin (*Pomc*) expression in the hypothalamus was dramatically induced by HFD feeding in both CO- and TCDD-exposed dams, but like leptin, *Pomc* levels were significantly lower in TCDDHFD dams compared to COHFD dams (**Figure 3F**). There was no effect of diet or chemical exposure on gene expression of the agouti-related peptide (*Agrp*), leptin receptor (*Lepr*), or insulin receptor (*Insr*) in the hypothalamus after 11 weeks of HFD feeding (**Figure 3F**). Neuropeptide Y (*Npy*) expression was decreased in TCDDChow dams compared to COChow dams, but unchanged in TCDDHFD dams (**Figure 3F**). Taken together, these data suggest that diet-induced weight gain in TCDD-exposed dams (relative to CO) may involve insufficient leptin levels, and consequently insufficient *Pomc* levels, thus promoting fat storage.

### 3.4. HFD feeding causes severe hyperglycemia in TCDD-exposed dams without altering insulin tolerance

Obesity is a major risk factor for developing diabetes, so although there were no long-term changes in fasting glycemia following TCDD exposure (**Figure 3A**), we investigated whether the diet-induced weight gain seen in TCDDHFD dams was associated with changes in glucose or insulin tolerance. As expected, COHFD dams displayed modest hyperglycemia during a glucose tolerance test (GTT) after 2.5 weeks of HFD feeding (**Figure 4A**). In contrast, TCDDHFD dams were severely hyperglycemic throughout the GTT, with a peak glycemia of ∼30 mM compared to ∼22 mM in COHFD dams (**Figure 4A**), and a significant increase in the overall glucose excursion compared to COHFD dams (**Figure 4A-AUC**). Following 6.5 (**Supplemental Fig. S4A**) and 9.5 weeks (**Figure 4C**) of HFD feeding, both COHFD and TCDDHFD dams had similar levels of hyperglycemia during a GTT. There were no differences in overall insulin tolerance between any groups at weeks 4, 7, or 10.5 of the metabolic challenge (**Figure 4B, 4D; Supplemental Fig. S4B**).

### 3.5. HFD feeding impairs insulin secretion in TCDD-exposed dams

Since defective glycemic control is often triggered by improper insulin secretion, we assessed changes in glucose-stimulated insulin secretion (GSIS) following HFD feeding. Corresponding with the profound hyperglycemia observed after 2.5 weeks of HFD feeding (**Figure 4A**), TCDDHFD dams had significantly lower plasma insulin levels 15 minutes following an i.p. glucose bolus compared to COHFD dams (**Figure 5Aiii**), and trended towards having reduced overall insulin secretion during the GTT compared to TCDDChow dams (**Figure 5Aiv**). Interestingly, after 6.5 weeks of HFD feeding, TCDDHFD dams had significantly higher basal insulin levels compared to COHFD dams (**Figure 5Biii**) and trended towards secreting more insulin overall during the GTT compared to TCDDChow dams (**Figure 5Biv**). After 9.5 weeks of HFD feeding, insulin levels in TCDDHFD dams had normalized back to COHFD levels (**Figure 5C iii,iv**). Prior TCDD exposure had no impact on insulin secretion in chow-fed dams (i.e. TCDDChow vs COChow) at any time point (**Figure 5Aii-5Cii**).

### 3.6. TCDDHFD dams have decreased islet size, a higher proportion of MAFA_-_ beta cells, and increased proinsulin accumulation compared to COHFD dams

Lastly, we assessed whether the observed changes in insulin secretion were associated with defects in islet morphology and other beta cell characteristics at week 11 of the metabolic challenge. As expected, HFD feeding caused a significant ∼2-fold increase in average islet size in control mice (**Figure 6A, 6H-J**). However, TCDDHFD dams did not show the same adaptation to HFD feeding, but rather showed an average islet size that was comparable to chow-fed dams (**Figure 6A, 6H-J**). TCDDHFD dams also had a significantly higher proportion of MAFA^-^ beta cells compared to all other groups (**Figure 6D, 6I**), suggesting a loss of beta cell identity. We also observed a significant increase in the proportion of beta cells with proinsulin accumulation throughout the cytoplasm as opposed to the expected perinuclear pattern of immunoreactivity in TCDDHFD mice compared to COHFD and TCDDChow dams **(Figure 6F, 6J**), although this was not associated with a change in circulating proinsulin:insulin ratio in cardiac blood from random-fed mice at week 11 of the metabolic challenge (**Figure 6G**). There was no change in % insulin^+^ area per islet (**Figure 6B, 6H**), % glucagon^+^ area per islet (**Figure 6C, 6H**), or % PCNA^+^ beta cells (**Fig. 6E**) in TCDDHFD dams compared to other groups. TCDDChow mice did not display any changes in islet composition compared to COChow dams (**Figure 6A-F, 6H-J**).

### 3.7. TCDDHFD females have reduced MafA, Nkx6.1 and Slc2a2 gene expression, and increased Pcsk1 expression

Due to our experimental design, we did not have sufficient animal numbers to isolate islets for RNA analysis from the dams in this study. Instead, we took advantage of RNAseq analysis on islets from a separate but related cohort of female mice. Briefly, non-pregnant female mice were injected with corn oil (CO; vehicle control) or 20 ng/kg/d TCDD 2x/week for 12 weeks, and were simultaneously fed either standard chow or 45% HFD (**Figure 7A**). Islets were isolated after 12 weeks of exposure, and gene expression was measured by RNAseq (**Figure 7B, 7D-E**) and qPCR (**Figure 7C**). To reiterate the main differences, the second model involved simultaneous exposure to TCDD and/or HFD in non-pregnant female mice (**Figure 7**), compared to the primary model of sequential exposure to TCDD transiently during pregnancy/lactation, followed by a 10-week period of chow feeding, then 11 weeks of chow or HFD feeding (**Figure 1-6**). Despite the differences, this second model provides a useful dataset to follow up on our histological observations (**Figure 6**).

Consistent with our histological finding of increased % MAFA^-^ beta cells in TCDDHFD dams (**Figure 6D, 6I**), RNAseq analysis showed that TCDDHFD females from the simultaneous exposure model had significantly lower *MafA* gene expression compared to COChow and TCDDChow females (**Figure 7B**). This finding was also validated by qPCR (**Figure 7C**). We next examined whether TCDD combined with HFD feeding altered the expression of other beta cell transcription factors. TCDDHFD females had significantly lower *Nkx6*.*1* expression compared to both TCDDChow and COHFD females (**Figure 7D**), supporting our hypothesis of beta cell dedifferentiation. Furthermore, we observed a diet-induced decrease in *Hnf4a*, irrespective of chemical exposure (**Figure 7D**), and a significant upregulation of *NeuroD1* in COHFD females compared to COChow only (**Figure 7D**). Next, we examined expression of genes involved in glucose-stimulated insulin secretion. Interestingly, TCDDHFD females had lower expression of the glucose transporter gene *Slc2a2* compared to TCDDChow females, suggesting a possible defect in glucose sensing (**Figure 7E**). The K_ATP_ channel regulatory subunit *Abcc8* was also downregulated in TCDDHFD females, however a similar decrease was observed in COHFD females and is therefore unlikely to contribute to a chemical-induced phenotype. Neither chemical or diet exposure affected the glucokinase gene *Gck*, or the K_ATP_ channel pore-forming subunit *Kcnj11* (**Figure 7E**). Finally, we assessed whether combined TCDD and HFD exposure affected proinsulin biosynthesis and/or processing by prohormone convertase enzymes PC1 and PC2 since we had observed a significant increase in proinsulin accumulation in beta cells from TCDDHFD dams in our study (**Figure 6F, 6J**). In the simultaneous exposure model, TCDDHFD females had a significant increase in *Pcsk1* expression compared to all other experimental groups, supportive of abnormal insulin processing in TCDDHFD conditions. There was no change in *Pcsk2*, and expression of *Ins1* and *Ins2* was affected by diet only (**Figure 7E**).

## 4. Discussion

Our study demonstrates that low-dose exposure to TCDD throughout pregnancy and lactation in mice has modest metabolic consequences during pregnancy but profoundly increases the susceptibility to weight gain and diabetes later in life. TCDD-exposed dams had normal body weight, basal blood glucose levels, plasma insulin levels, and beta cell plasticity during pregnancy, but were transiently hypoglycemic at birth and heavier at 2 weeks post-birth compared to CO-exposed dams. After chemical exposure stopped at the end of lactation, TCDD-exposed dams maintained normal glucose homeostasis, but showed a modest increase in body weight compared to CO-exposed dams within 5 weeks post-TCDD. Once challenged with HFD feeding, TCDD-exposed dams rapidly became obese, severely glucose intolerant, and showed beta cell dysfunction. The TCDD-induced exacerbation of glucose intolerance and dysregulation of insulin secretion eventually resolved, but TCDDHFD mice remained significantly heavier than COHFD mice throughout the study. TCDDHFD dams also displayed defects in insulin processing, reduced islet size, and a larger proportion of MAFA^-^ beta cells compared to COHFD dams. Taken together, these data suggest that low-dose dioxin exposure in female mice leads to sustained susceptibility to developing obesity and diabetes long after exposure ceases.

To our knowledge, this is the first study assessing the association between chronic low-dose dioxin exposure during pregnancy/lactation and GDM or long-term diabetes risk in a mouse model. Our study shows that dioxin dramatically induces *Cyp1a1* expression in the pancreas during pregnancy, which is indicative of direct chemical exposure to pancreas tissue. Interestingly, unlike our previous studies in non-pregnant mice [8,9], the degree of *Cyp1a1* induction in the pancreas was much higher than in the liver mid-pregnancy. It is unclear whether pregnancy promotes greater dioxin accumulation in the pancreas compared to other tissues or if the pancreas may be more sensitive to AhR-mediated dioxin signaling mid-pregnancy. We also observed a decrease in *AhR* and *Arnt* expression in the pancreas but not liver mid-pregnancy, indicating negative feedback regulation of the AhR pathway. Interestingly, we previously reported upregulation of Cyp1a1 enzyme activity for at least 2 weeks in islets following acute TCDD exposure [8], suggesting that Cyp1a1 activation may persist after AhR suppression. Whether induction of *Cyp1a1* or suppression of *AhR* in the pancreas following chronic low-dose dioxin exposure has protective or deleterious effects remains an important area of investigation.

Based on our previous work with acute high-dose TCDD [8,9], we predicted that chronic low-dose TCDD exposure to pancreas tissue during pregnancy could have both immediate and long-term implications on islet physiology and diabetes risk. Interestingly, low-dose TCDD exposure alone did not cause overt GDM in this study. In fact, rather than developing hyperglycemia, we found that TCDD-exposed dams were transiently hypoglycemic at birth. Labour is an acute form of metabolic stress characterized by elevated blood glucose levels to compensate for increases in energy demand [31]. TCDD-induced hypoglycemia at birth suggests that dioxin exposure during pregnancy may impair the ability to adapt to a metabolic stressor; however, in the absence of a prolonged stressor the dams return to euglycemia. Future studies should investigate whether combining TCDD exposure during pregnancy with a secondary metabolic stressor, such as HFD feeding, would trigger the development of GDM.

Thus far, epidemiological studies investigating the association between POP exposure during pregnancy and diabetes incidence have been inconsistent. Some studies reported a positive association between GDM and serum POP levels, including polychlorinated biphenyls (PCB) [22– 24], polyfluoroalkyl (PFAS) [23], and polybrominated diphenyl ethers (PBDEs) [23,25]. A positive association between GDM and self-reported pesticide exposure during pregnancy has also been reported [32]. However, other studies showed no association between GDM incidence and serum organochloride pesticides [25,26] or PCB levels [26,27]. Inconsistencies between human studies can result from the inability to control confounding variables, such as varying POP concentrations between participants, differences in timing of POP measurements, method of diabetes diagnosis, genetic variation, and environmental factors. Using a controlled mouse model, we demonstrated that chronic low-dose TCDD exposure did not impair body weight, basal blood glucose, plasma insulin levels, or beta cell plasticity acutely during pregnancy, but rather caused long-term metabolic dysregulation. It is important to note that we did not have non-pregnant controls in this study and therefore we cannot determine whether the long-term effects of TCDD on metabolism were influenced by the timing of exposure during pregnancy/lactation. Future studies should include a head-to-head comparison of the effects of TCDD on metabolism in pregnant versus non-pregnant females. Regardless, our study still highlights the need to track the metabolic health of mothers long-term following pollutant exposure during pregnancy.

The most persistent consequence of transient low-dose TCDD exposure during pregnancy and lactation in our model was excessive weight gain later in life. TCDD-exposed dams showed modest weight gain in the absence of a metabolic stressor, but rapidly developed pronounced obesity following a transition to HFD feeding. These results are interesting given that acute high-dose TCDD exposure is known to cause wasting syndrome [8,33]. To our knowledge this is the first study to report an increase in weight gain following chronic low-dose TCDD exposure. Whether low-dose TCDD exposure also promotes weight gain in non-pregnant female mice remains to be investigated. Obesity in TCDDHFD dams was also associated with abnormal energy homeostasis. TCDDHFD dams had significantly reduced circulating leptin levels and hypothalamic *Pomc* levels compared to COHFD dams. Leptin is essential for regulating energy homeostasis and can activate POMC neurons in the hypothalamus [34,35]. *Pomc* expression, especially in LepR POMC neurons, is critical for maintaining normal energy balance by providing anorexigenic drive [36]. Both leptin [37] and melanocortins produced by POMC neurons [38] can directly regulate peripheral lipid metabolism, and in effect leptin- and POMC-deficiency cause obesity by increasing food intake, limiting energy expenditure, and promoting fat storage [39–41]. We did not observe significant changes in food intake between TCDDHFD and COHFD dams but did find that TCDDHFD dams had significantly higher % fat mass than COHFD mice, suggesting an adaptation in energy expenditure and/or energy storage that promotes obesity. Studies using metabolic chambers would provide additional insight into the mechanisms underlying TCDD-induced weight gain. Regardless, our mouse model suggests that transient low-dose TCDD exposure during pregnancy and lactation may contribute to obesity pathogenesis.

Transient TCDD exposure also accentuated the development of diet-induced hyperglycemia, which appears to be primarily driven by impaired beta cell plasticity and function. The rapid onset of hyperglycemia in TCDD-exposed mice after 2.5 weeks of HFD feeding coincided with suppressed glucose-induced insulin secretion *in vivo*, but no change in peripheral insulin sensitivity. Interestingly, the hypoinsulinemia in TCDDHFD dams transitioned to hyperinsulinemia, before eventually re-equilibrating back to COHFD levels, pointing to an initial defect in insulin secretion at the beta cell level. Although TCDD-induced exacerbation of glucose intolerance and dysregulation of insulin secretion eventually resolved, TCDDHFD dams showed lasting defects in islet morphology, beta cell maturity, and proinsulin processing that could have long-term implications on metabolism. TCDDHFD dams had decreased islet size compared to COHFD dams, suggesting that TCDD-exposed beta cells lack the adaptive proliferative response to the metabolic demand of HFD feeding. However, without tissues from different timepoints, we cannot rule out the possibility that TCDDHFD dams experienced an appropriate increase in beta cell proliferation earlier, followed by an increase in beta cell apoptosis. We also found that TCDDHFD dams had a modest but significant increase in MAFA^-^ beta cells, which was validated at the gene level in a non-pregnancy mouse model with simultaneous TCDD and HFD exposure. MAFA is a transcription factor essential for maintaining beta cell maturity and function. Reduced *MAFA* expression is associated with beta cell dedifferentiation and beta cell dysfunction in diabetes progression [42,43]; MAFA is also commonly supressed in beta cells from humans with type 2 diabetes [44,45]. Additionally, we observed a decrease in *Nkx6*.*1* and *Slc2a2* in TCDDHFD females in the simultaneous exposure model, which is characteristic of immature beta cells [46– 49]. Lastly, TCDDHFD dams also had increased proinsulin accumulation, indicating impaired proinsulin synthesis and/or processing. Interestingly, in the non-pregnancy model TCDDHFD islets had no change in *Ins1* or *Ins2* but increased *Pcsk1* expression compared to COHFD islets, potentially as an adaptive response to increased proinsulin accumulation. Taken together, these results suggest that TCDD exposure may induce beta cell dedifferentiation following HFD feeding, which contributes to increased diabetes risk.

Epidemiological studies consistently show an association between exposure to dioxin/dioxin-like compounds and diabetes incidence, but limited studies examine dioxin exposure during pregnancy. Our study shows that chronic low-dose TCDD exposure during pregnancy was insufficient to induce GDM, but eventually led to impaired metabolic adaptability to HFD feeding, including glucose intolerance, beta cell dysfunction, and increased weight gain in mice. Our study suggests that background exposure to dioxin or dioxin-like chemicals could be contributing to obesity and diabetes incidence. The effects of low-dose pollutant exposure on metabolism, particularly in females, should be further investigated.

## Abbreviations

AhR: Aryl hydrocarbon receptor
CO: Corn Oil
Cyp: Cytochrome P450
Cyp1a1: Cytochrome P450 1A1
Cyp1a2: Cytochrome P450 1A2
GDM: Gestational diabetes mellitus
GSIS: Glucose stimulated insulin secretion
GTT: Glucose tolerance test
HFD: High fat diet
ITT: Insulin tolerance test
POPs: Persistent Organic Pollutants
TCDD: 2,3,7,8-tetrachlorodibenzo-*p*-dioxin

## 5. Acknowledgements

We thank the Genomics Laboratory at Health Canada for their help conducting RNAseq analysis. This research was supported by a Canadian Institutes of Health Research (CIHR) Project Grant (#PJT-2018-159590), Natural Sciences and Engineering Research Council of Canada (NSERC) Discovery Grant (RGPIN-2017-06272), the Canadian Foundation for Innovation John R. Evans Leaders Fund (#37231), and an Ontario Research Fund award. M.P.H. was supported by an Ontario Graduate Scholarship (OGS) and CIHR CGS-D award. M.A.P. was supported by an OGS and Queen Elizabeth II Scholarship. H.B. received a NSERC Undergraduate Student Research Award (USRA) and a Carleton University Walker Summer Research Award. C.D.S. received a NSERC USRA and Carleton University I-CUREUS internship award. J.E.B. is the guarantor for this work.

## 6. Author Contributions

J.E.B. and M.P.H. conceived the experimental design and wrote the manuscript. M.P.H., R.C.M., H.B., C.D.S., M.A.P., D.I.M.A., M.Z., G.M., M.J.C., and J.E.B. were involved with acquisition, analysis, and interpretation of data. All authors contributed to manuscript revisions and approved the final version of the article.

## 7. Declaration of Interest

The authors declare no competing interests.

## Notes

### Competing Interest Statement

The authors have declared no competing interest.

### Summary of Updates

RNAseq data from a separate but related cohort of non-pregnant female mice has been added to this paper to support histological findings in our pregnant mice.

