## Supplemental Data for "Female mice exposed to low-doses of dioxin during pregnancy and lactation have increased susceptibility to diet-induced obesity and diabetes"

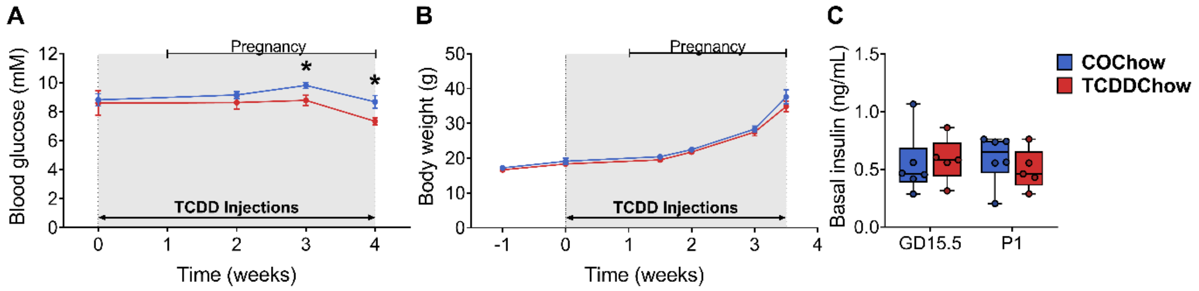


**Fig. S1: Chronic low-dose TCDD exposure during pregnancy causes hypoglycemia at birth.** A second cohort of female mice were injected with either corn oil or 20 ng/kg/d TCDD 2x/week during mating and pregnancy and euthanized at postnatal day 1 (P1) (see **Figure 1A** for study timeline). (**A**) Blood glucose and (**B**) body weight were measured weekly following a 4-hour morning fast. (**C**) Plasma insulin levels were measured at gestational day (GD) 15.5 and P1 following a 4-hour morning fast. All data are presented as mean ± SEM in line graphs or median with min/max values in box and whisker plots. Individual data points on box and whisker plots represent biological replicates (different mice). *p <0.05 versus control. The following statistical tests were used: (**A-B**) two-way REML-ANOVA with uncorrected Fisher's LSD test; (**C**) two-tailed unpaired t-test.

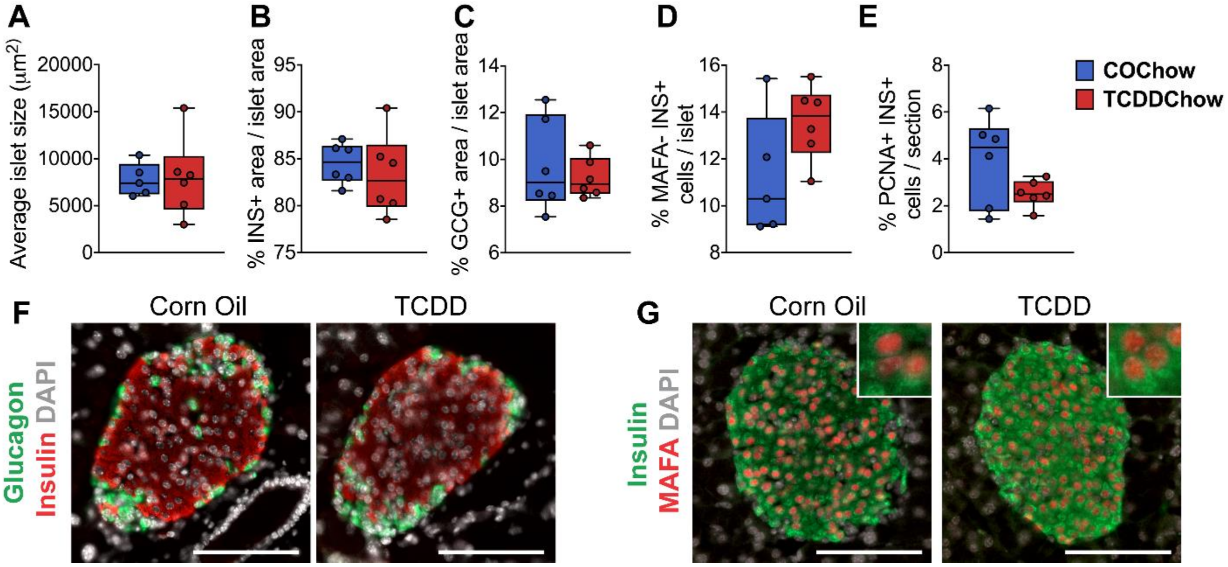


**Fig. S2: Chronic low-dose TCDD exposure during pregnancy did not alter islet morphology at birth.** Whole pancreas tissue was harvested from dams at postnatal day 1 for analysis by immunofluorescence staining (see **Figure 1A** for study timeline). (**A**) Average islet area. (**B-C**) % of islet area that is immunoreactive for (**B**) insulin, or (**C**) glucagon. (**D-E**) % of insulin$^{+}$ cells per islet that are (**D**) MAFA$^{-}$, or (**E**) PCNA$^{+}$. (**F-G**) Representative images of pancreas sections showing immunofluorescence staining for (**F**) insulin/glucagon or (**G**) insulin/MAFA. Inset regions in (**G**) show MAFA$^{+}$INS$^{+}$ cells. Scale bar = 100 µm. All data are presented as median with min/max values. Individual data points on box and whisker plots represent biological replicates (different mice). The following statistical tests were used: (**A-D**) two-tailed unpaired t-test; (**E**) two-tailed Mann Whitney Test.

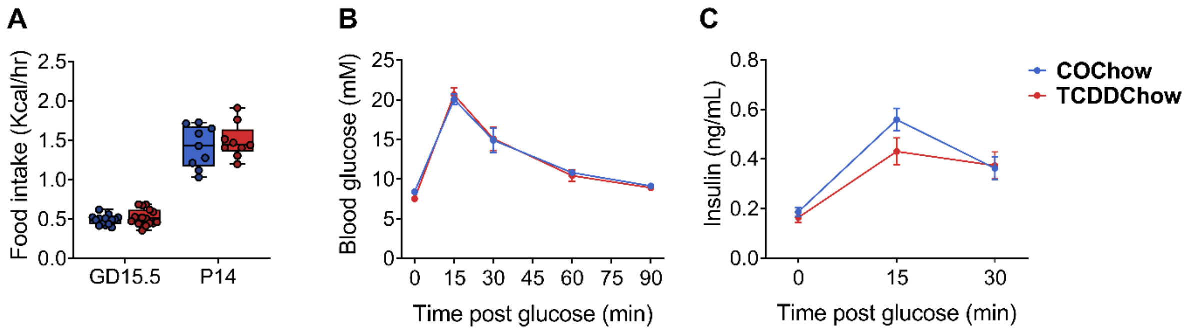


**Fig. S3: TCDD exposure during pregnancy did not have any acute effects on food intake or long-term effects on metabolism.** Female mice were injected with either corn oil of 20 ng/kg/g TCDD during mating, pregnancy, and lactation, and then tracked up to 10 weeks (see **Figure 1A** for study timeline). (**A**) Caloric intake was measured at gestational day (GD) 15.5 and mid-lactation on postnatal day (P) 14. (**B**) Blood glucose and (**C**) plasma insulin levels during a glucose tolerance test at 3 weeks post-TCDD (i.e. week 10 of the study). All data are presented as mean ± SEM in line graphs or median with min/max values in box and whisker plots. Individual data points on box and whisker plots represent biological replicates (different mice). The following statistical tests were used: (**A**) two-tailed unpaired t-test; (**B**) two-way RM-ANOVA with Sidak test; (**C**) two-way REML ANOVA with Sidak test.

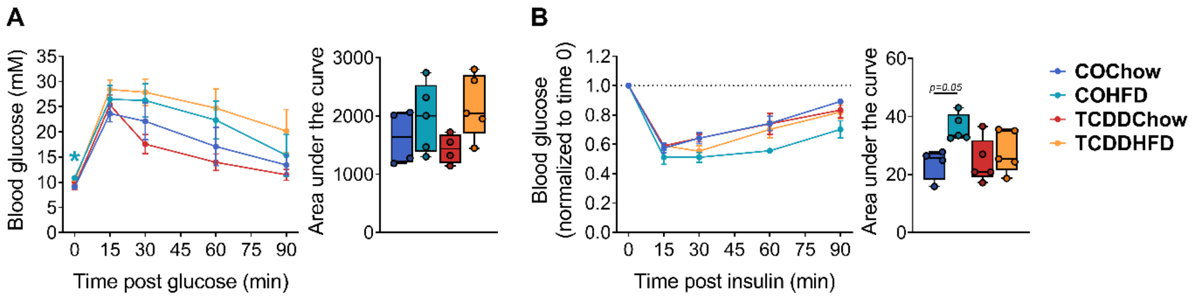


**Fig. S4: TCDD-exposed dams had normal glucose and insulin tolerance after 6.5-7 weeks of HFD feeding.** Glucose tolerance (GTT) and insulin tolerance (ITT) were assessed *in vivo* at week 6.5-7 of the metabolic challenge (see **Figure 1A** for study timeline). (**A**) Blood glucose levels during a GTT at week 6.5 of the metabolic challenge. (**B**) Blood glucose levels during an ITT at week 7 of the metabolic challenge. All data are presented as mean ± SEM in line graphs or median with min/max values in box and whisker plots. Individual data points on box and whisker plots represent biological replicates (different mice). The following statistical tests were used: (**A-B**) line graph, two-way RM ANOVA with Tukey multiple comparison test; box and whisker plots, two-way ANOVA with Tukey's multiple comparison test.

**Supp. Table 1:** qPCR primer sequences.

| **Target** | **Forward Primer** | **Reverse Primer** |
|---|---|---|
| *Agrp* | TGT AAG GCT GCA CGA GTC C | TTG AAG AAG CGG CAG TAG CAC |
| *Ahr* | AGC CGG TGC AGA AAA CAG TAA | AGG CGG TCT AAC TCT GTG TTC |
| *Arnt* | GAC AGA CCA CAG GAC AGT TCC | AGC ATG GAC AGC ATT TCT TGA A |
| *Cyp1a1* | ATC ACA GAC AGC CTC ATT GAG C | AGA TAG CAG TTG TGA CTG TGT C |
| *Cypa* | GGT GGA GAG CAC CAA GAC AGA | GCC GGA GTC GAC AAT GAT G |
| *Hprt* | GCT GAC CTG CTG GAT TAC AT | TTG GGG CTG TAC TGC TTA AC |
| *Insr* | AAA TGC AGG AAC TCT CGG AAG CCT | ACC TTC GAG GAT TTG GCA GAC CTT |
| *Lepr* | TGG ATG AAA GGG GAC TTG AC | GGC ACA TGA CAT TCA CAT CC |
| *Mafa* | AGT CGT GCC GCT TCA AG | CGC CAA CTT CTC GTA TTT CTC C |
| *Npy* | CCC CAG AAC AAG GCT TGA AG | TTG GAA AAG TCG GGA GAA CAA |
| *Pomc* | TGG GTC ACTT CCG CTG GG | TCC TCC GCA CGC CTC TG |
| *Ppia* | AGC TCT GAG CAC TGG AGA GA | GCC AGG ACC TGT ATG CTT TA |

**Supp. Table 2:** Summary of statistical tests used and verification of parametric assumptions.

| Figure | Stats Test Used | Passed Variance? | Passed Normality? |
|---|---|---|---|
| 1B | Two-tailed unpaired T-test for pancreas, liver, and fat. Two-tailed Mann Whitney test for placenta | All tissues passed except placenta | Yes |
| 1C-D | Two-tailed unpaired T-test | Yes | Yes |
| 1E-I | Two-tailed unpaired T-test | Yes | Yes |
| 2A: Line Graph | Two-way REML ANOVA | N/A | N/A |
| 2A: Inset Graph | Two-tailed unpaired T-test | Yes | Yes |
| 2B | Two-way REML ANOVA | N/A | N/A |
| 2C | Two-tailed Mann Whitney test at GD15.5, P0, and P21. Two-tailed unpaired t-test at P14 | GD15.5 and P0 failed. P14 and P21 passed | GD15.5, P0, and P21 failed. P14 passed |
| 2D | Two-way RM ANOVA | Yes | Yes |
| 2E | Two-way REML ANOVA | Yes | Yes |
| 2F-H | Two-way RM ANOVA | Yes | Yes |
| 3A: Line Graph | Two-way RM ANOVA | Yes | TCDDChow failed |
| 3A: AUC | Two-way ANOVA | Yes | TCDDChow failed |
| 3B: Line Graph | Two-way RM ANOVA | Yes | Yes |
| 3B:AUC | Two-way ANOVA | Yes | Yes |
| 3C | Two-way ANOVA | Yes | No |
| 3D-E | Two-way ANOVA | Yes | Yes |
| 3F | Two-way ANOVA | TCDDChow group failed for NPY, InsR, and POMC targets | Yes |
| 4A: Line Graph | Two-way RM ANOVA | Yes | Yes |
| 4A: AUC | Two-way ANOVA | Yes | Yes |
| 4B: Line Graph | Two-way RM ANOVA | TCDDHFD failed | Yes |
| 4B: AUC | Two-way ANOVA | TCDDHFD failed | Yes |
| 4C: Line Graph | Two-way RM ANOVA | Yes | Yes |
| 4C: AUC | Two-way ANOVA | Yes | Yes |
| 4D: Line Graph | Two-way RM ANOVA | Yes | Yes |
| 4D: AUC | Two-way ANOVA | Yes | Yes |
| 5A: Line Graphs | Two-way RM ANOVA | Yes | Yes |
| 5A: AUC | Two-way ANOVA | Yes | Yes |
| 5B: Line Graphs | Two-way RM ANOVA | Yes | COHFD failed |
| 5B: AUC | Two-way ANOVA | Yes | COHFD failed |
| 5C: Line Graphs | Two-way RM ANOVA | Yes | Yes |
| 5C: AUC | Two-way ANOVA | Yes | Yes |
| 6A-D | Two-way ANOVA | Yes | Yes |

| | | | |
|---|---|---|---|
| 6E | Two-way ANOVA | Yes | COHFD, TCDDChow, and TCDDHFD failed |
| 6F-G | Two-way ANOVA | Yes | Yes |
| 7B | Two-way ANOVA | Yes | TCDDHFD failed |
| 7C | Two-way ANOVA | Yes | Yes |
| 7D | Two-way ANOVA | Yes | *Hnf4a* COChow failed |
| 7E | Two-way ANOVA | Yes | *Pcsk1* TCDDChow failed |
| S1A-B | Two-way REML ANOVA | Yes | Yes |
| S1C | Two-tailed unpaired T-test | Yes | Yes |
| S2A-D | Two-tailed unpaired T-test | Yes | Yes |
| S2E | Two-tailed Mann Whitney Test | No | Yes |
| S3A | Two-tailed unpaired T-test | Yes | Yes |
| S3B | Two-way RM ANOVA | Yes | Yes |
| S3C | Two-way REML ANOVA | Yes | Yes |
| S4A-B | Two-way RM ANOVA | Yes | Yes |
